# Bayesian inference of population structure using identity-by-descent-based stochastic block models

**DOI:** 10.1101/2025.11.26.690810

**Authors:** Steven J. Micheletti, Katarzyna Bryc, Samantha G. Ancona Esselmann, Peter R. Wilton, the 23andMe Research Team, William A. Freyman

**Affiliations:** 23andMe Research Institute, Palo Alto CA, USA; Darwin’s Ark, Broad Institute of MIT and Harvard, Cambridge, MA, USA

## Abstract

Fine-scale population structure is increasingly studied by clustering identity-by-descent (IBD) haplotypes, yet most current approaches rely on heuristic, modularity-based algorithms that can over-partition noisy IBD graphs and provide no explicit measure of uncertainty. We introduce a fully Bayesian framework that models IBD sharing with a generative planted-partition stochastic block model (PPSBM). To benchmark accuracy, we simulated genomes under recent population divergence and compared PPSBM estimates with those from the widely used Leiden community-detection algorithm. The PPSBM correctly assigned 81.0% of individuals on average versus 67.0% for Leiden, outperforming Leiden in 92.0% of replicates. Posterior probabilities from the PPSBM reflected patterns of recent shared ancestry or admixture, whereas Leiden tended to merge such clusters or assign individuals deterministically. Furthermore, we applied the method to the genomes of 63,196 individuals to reveal fine-scale population structure in Mexico, including multiple indigenous communities and diasporic groups such as Lebanese Mexicans and Syrian Jewish Mexicans. Our results demonstrate that a probabilistic, IBD-based PPSBM yields more accurate and biologically interpretable population assignments than popular heuristic methods, while simultaneously quantifying uncertainty and accommodating admixed genomes. The method scales to thousands of individuals and provides a principled foundation for downstream demographic inference and association studies in the presence of subtle structure.

## 1 Introduction

Understanding population structure is essential for studying genetic diversity, inferring evolutionary history, and avoiding confounding in association studies (Price et al. 2006; Jakobsson et al. 2008; Li et al. 2008). Population structure reflects the genetic relationships among individuals, shaped by ancestry, migration, and demographic events, and can be complex—especially since genetic variation exists along gradients rather than in discrete groups. Several complementary approaches have been developed to examine population structure. Principal component analysis (PCA) is often used to identify major axes of genetic variation using single-nucleotide polymorphism (SNP) data, and can help visualize genetic structure and investigate the spatial distribution of genetic variation (Novembre et al. 2008). Many model-based approaches also rely on SNP allele frequencies; for example, STRUCTURE clusters individuals into one or more latent populations, explicitly modeling admixture (Pritchard et al. 2000). Approaches for detecting population structure that take advantage of haplotype data are a powerful alternative to those based on SNP data (Browning and Weir 2010). By leveraging patterns of linkage disequilibrium (LD) and haplotype variation, these methods may be less biased by SNP ascertainment (Conrad et al. 2006).

Recently, studies have used clustering of identical-by-descent (IBD) haplotypes to identify fine-scale population structure (Nait Saada et al. 2020; Dai et al. 2020; Han et al. 2017). IBD refers to segments of the genome that are shared between individuals due to inheritance from a common ancestor. Clustering individuals based on patterns of IBD sharing enables the detection of more recent and subtle structure, since long shared haplotypes reflect very recent genetic relatedness that may be missed by SNP allele frequency-based methods. Non-probabilistic clustering algorithms, such as hierarchical agglomerative clustering or the Louvain method (Blondel et al. 2008), are typically used for this task due to their scalability and simplicity. However, heuristic clustering methods like Louvain that maximize modularity (Newman and Girvan 2004) are prone to overfitting the data by detecting spurious structure, especially when the data is noisy or sparse (Peixoto 2023).

Here, we demonstrate that a probabilistic approach to clustering IBD haplotype data can accurately resolve fine-scale population structure. By applying stochastic block models (SBMs; Holland et al. 1983) with Bayesian inference to cluster IBD segment data, our method provides a principled framework for modeling uncertainty and capturing admixture, allowing individuals to belong to multiple genetic groups with varying degrees of membership.

## 2 Methods

SBMs are probabilistic generative models for graphs (Holland et al. 1983). They are a highly flexible approach for modeling the structure of a graph. Here we use a special case of the general SBM called the planted partition SBM (PPSBM), which identifies assortative structure within a graph (Condon and Karp 2001). Assortative structure is present when the graph can be partitioned into groups in which vertices of the same group are more likely to be connected by an edge than vertices of different groups.

We are interested in identifying population structure within large sets of genetic data. To do this, we first estimate the pairwise amount of IBD shared between all individuals in the dataset. The IBD may be computed using partial genomes, in which regions of the genome are masked out so that IBD segments cannot be detected within them if they are not assigned to a specific population by a local ancestry inference algorithm. This is useful when some individuals in the dataset are admixed, and we wish to detect populations structured through specific ancestry components. These IBD estimates are used to construct the IBD graph *A*, in which each vertex represents an individual and the number of edges *A*_*ij*_ between vertices *i* and *j* is calculated as:

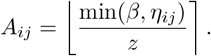

In this equation, *η*_*ij*_ is the amount of IBD shared between individuals *i* and *j, z* is a divisor that adjusts how the pairwise IBD is weighted in the IBD graph, and *β* is the maximum amount of IBD that will be considered for a pairwise relationship. *A*_*ii*_ represents self IBD or runs of homozygosity (ROH). In practice, we typically ignore all ROH.

We wish to infer assortative structure within the IBD graph in which genetic groups (*i*.*e*., populations) consist of individuals who share more IBD with one another than to individuals in other groups. Using Bayesian inference, we infer *b*, the vector of assignments of individuals to genetic groups, given the observed IBD graph *A*):

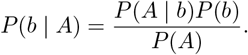

Following Zhang and Peixoto (2020), the likelihood for the PPSBM of observing the IBD graph given the assignment of individuals to genetic groups is given by:

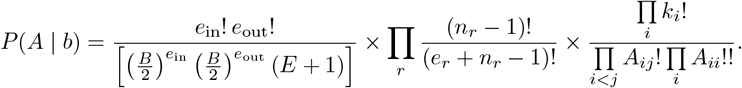

In this likelihood, *B* is the number of genetic groups, *E* is the total number of edges in *A, e*_in_ is the total number of edges within genetic groups, *e*_out_ is the total number of edges between different genetic groups, *e*_*r*_ is the number of edges for genetic group *r, n*_*r*_ is the number of individuals in genetic group *r*, and *k*_*i*_ is the degree of individual *i*. To perform inference we also need to calculate the prior probability on the assignment of individuals to genetic groups, where *N* is the total number of individuals:

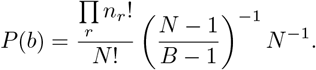

See Peixoto (2019) for a derivation of this prior distribution.

We estimate the posterior distribution of genetic group assignments by sampling group assignments using Markov chain Monte Carlo (MCMC; Metropolis et al. 1953), which enables us to avoid computing the intractable normalization term *P* (*A*). The MCMC algorithm uses merge-split proposals as described in Zhang and Peixoto (2020). To summarize the posterior distribution, we align the labels of the clustering partitions across MCMC samples using a modified form of the method described in Peixoto (2021). In practice, an MCMC initialized with a random number of genetic groups and individuals randomly assigned to those groups may be slow to converge. In these cases, we combine samples from multiple independent MCMCs, where each MCMC is initialized with an estimate of the genetic group assignments that maximizes modularity.

## 3 Simulation study

To simulate patterns of IBD sharing under recent population divergence, we used the coalescent simulator `msprime`(Baumdicker et al. 2022). We set up a simple hierarchical demographic scenario involving six derived populations (A–F) which split from two intermediate ancestral populations (ABC and DEF) four generations in the past. ABC and DEF, in turn, split from a single ancestral population (ABCDEF) eight generations in the past (Figure 1). The effective population size of ABCDEF was 1,000, and each derived population evenly split the size of the ancestral population. We simulated ancestry over a 249 Mb region (approximating human chromosome 1) with a uniform recombination rate of 1e−8 per base pair per generation. We sampled 20 diploid individuals from each of the six derived populations. To quantify IBD sharing, we extracted all pairwise IBD segments from the simulated tree sequence with length ≥ 5 Mb using `tskit`(Kelleher et al. 2016; Wong et al. 2024). For each individual pair, we summed the total length of shared IBD segments.

**Figure 1:**
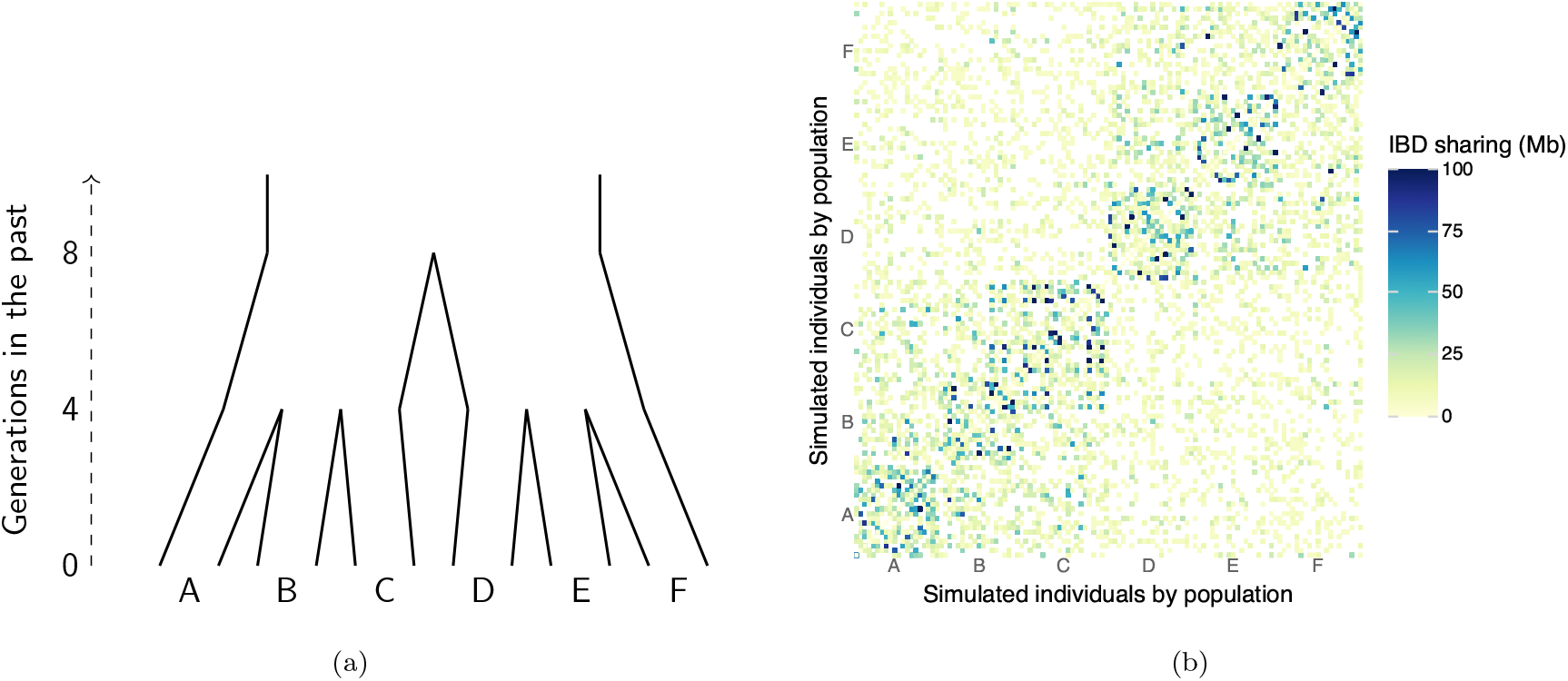
(a) Simulating recent and fine-scale population structure. Populations A-C and D–F split from ABC and DEF respectively at 4 generations in the past, and ABC and DEF split from ABCDEF at 8 generations in the past. **(b) IBD shared between simulated individuals for a single simulation replicate**. Chromosomes were simulated for twenty individuals from each of the six populations A-F. See the main text for details about the simulations. Cells are colored by the pairwise amount of IBD shared in megabases.

For 1000 simulation replicates, pairwise IBD sharing was used to infer population structure with both the PPSBM as described above and the Leiden algorithm. The Leiden algorithm is an improved version of the Louvain algorithm that attempts to find the partition of the graph that maximizes modularity (Traag et al. 2019). We used the PPSBM as implemented in the `graph-tool`Python library (Peixoto 2014) and the Leiden algorithm as implemented in `leidenalg`(Traag 2023). To compare estimated PPSBM population assignments with the true simulated populations, we resolved label ambiguity by aligning each estimated population with the true labels based on maximum overlap. Specifically, for individuals simulated from population A, we identified the most frequently assigned PPSBM population and relabeled it as population A. This majority-vote approach was applied to each true population to establish a consistent mapping between estimated and true population labels. We repeated this process for the Leiden population assignments. Next, we determined the accuracy of each method by counting the number of individuals in each simulation replicate that were correctly assigned to their true simulated population. For the PPSBM, we considered the maximum a posteriori (MAP) assignment of individuals to populations.

In 920 out of 1000 simulation replicates, the MAP PPSBM population assignments were more accurate than the Leiden algorithm (Figure 2a). On average across all replicates, the PPSBM accurately assigned 14.0% more individuals to the correct population compared to the Leiden algorithm. On average, the PPSBM assigned 81.0% of individuals to the correct population, while Leiden was 67.0% accurate on average. Figure 2b shows the results for a single representative simulation replicate. In this replicate, the MAP PPSBM population assignments were correct for 90.0% of individuals, and the Leiden population assignments were correct for 76.7% of individuals. This example illustrates the different error modes of the two approaches. For this simulation replicate both methods struggled to distinguish individuals from recently diverged populations E and F. The PPSBM handled this uncertainty through lower posterior probabilities for each population assignment. The Leiden algorithm, on the other hand, simply lumped the two populations together. The posterior probabilities of the PPSBM could be interpreted biologically as a signal of admixture or recent shared ancestry. For example, the individuals in populations E and F who get significant non-zero posterior probabilities for H make up a subpopulation of individuals who have closer genetic connections across E and F than the other individuals in E and F. By allowing individuals to belong to multiple populations, the PPSBM can capture signals of admixture and genetic variation that exists along gradients; an individual is not forced into a single population.

**Figure 2:**
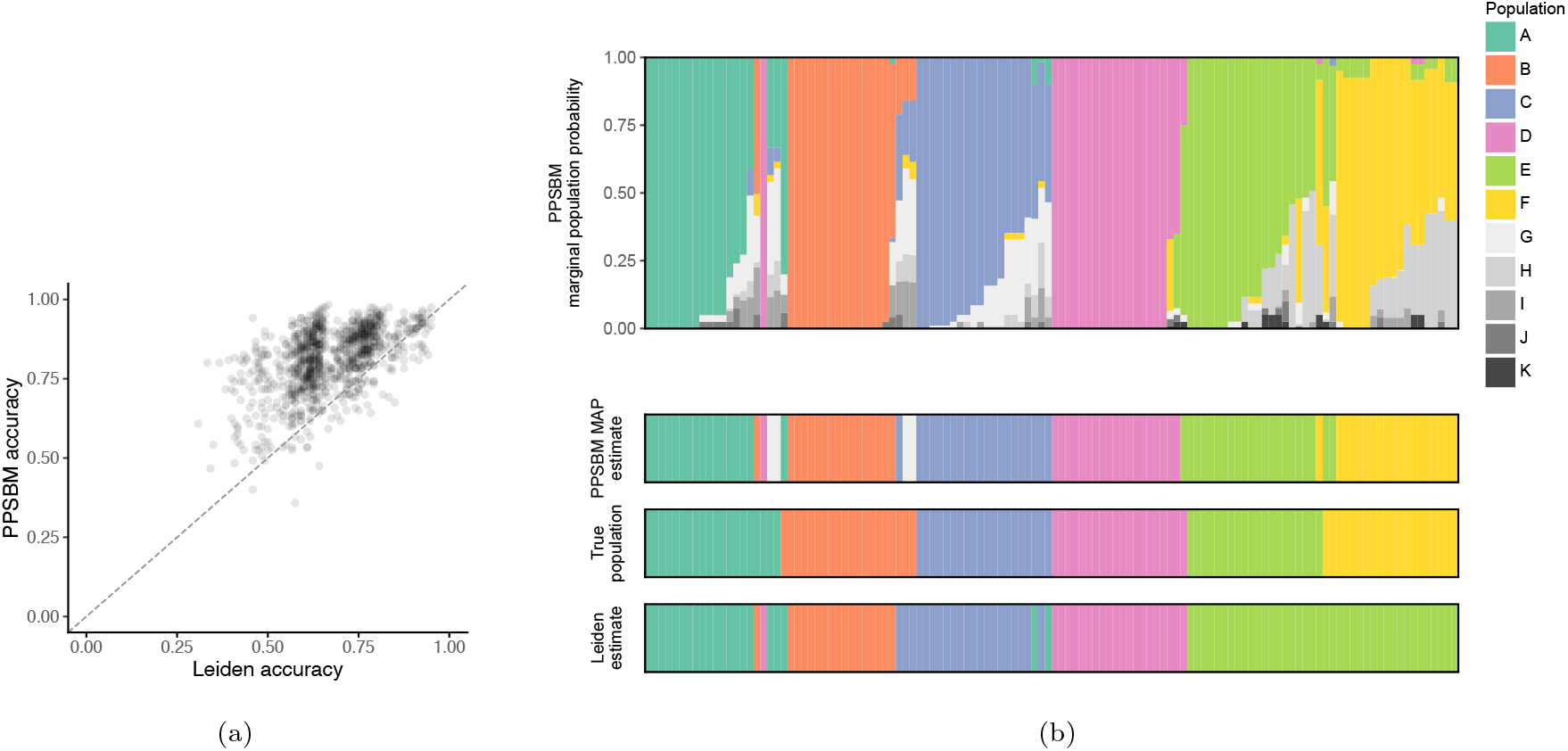
(a) Accuracy of the PPSBM compared to the Leiden algorithm in 1000 simulation replicatesx. The PPSBM assigned more individuals to the correct population than the Leiden algorithm in 92% of simulation replicates. On average, the PPSBM assigned 81.0% of individuals to the correct population on average, while Leiden was 67.0% accurate. **(b) Comparison of PPSBM and Leiden estimates for a representative simulation replicate**. The top panel shows the marginal population probabilities from the PPSBM for a single simulation replicate, where each column represents an individual. The next panels down show (1) the MAP assignments of individuals to populations using the PPSBM, (2) the true populations A, B, C, D, E, and F simulated under the population history shown in Figure 1, and (3) the Leiden estimates. Each column represents the same simulated individual in all panels. The MAP PPSBM assignments were correct for 90.0% of individuals, and the Leiden population assignments were correct for 76.7% of individuals. See the main text for a discussion on these results.

## 4 Empirical application

To demonstrate the ability of IBD-based PPSBM clustering to identify granular population structure, we applied the method to the IBD shared among 63,196 research-consented participants in the 23andMe database who reported that all four of their grandparents were born in Mexico (Figure 3).

**Figure 3:**
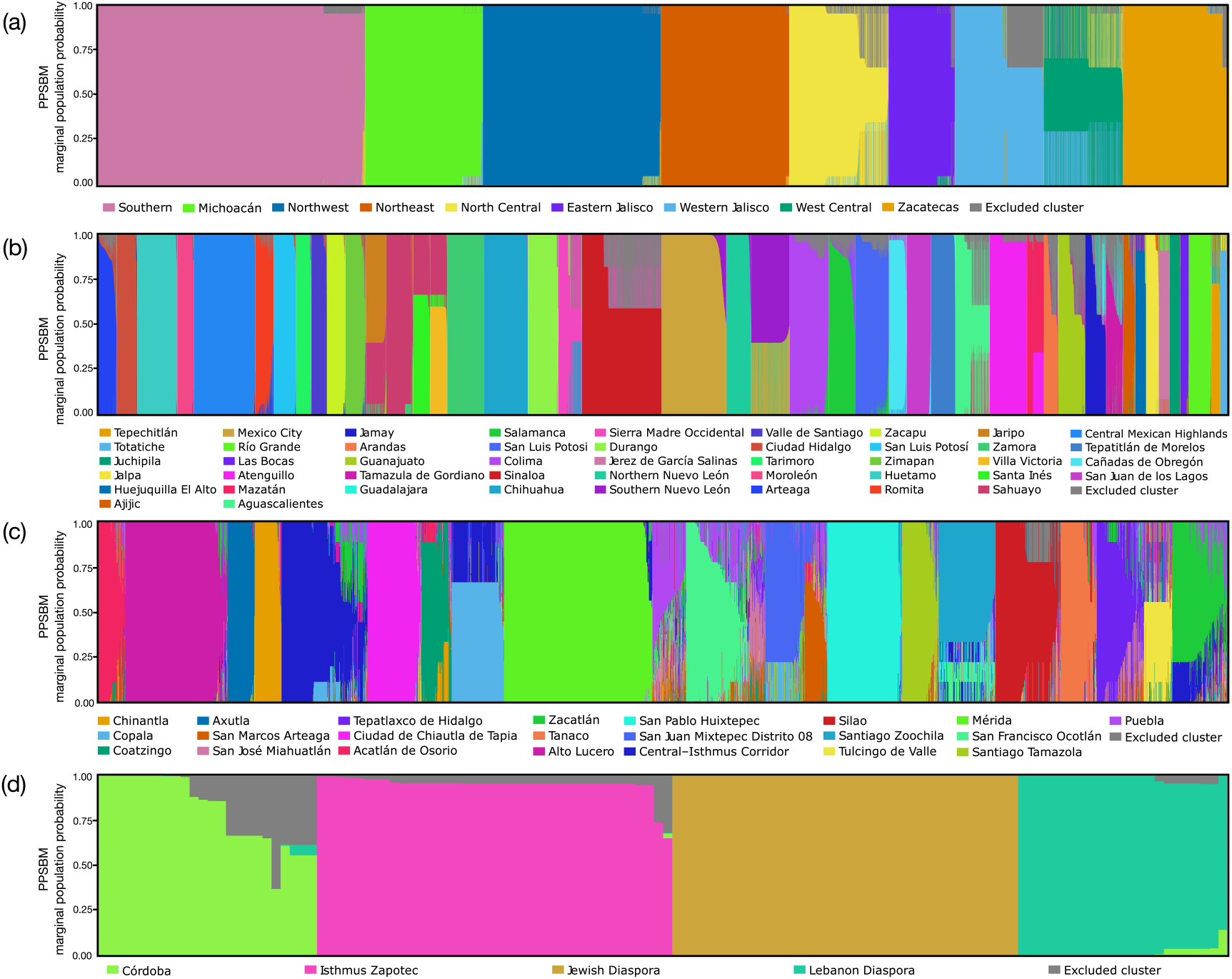
Marginal assignment probabilities of individuals with all four grandparents born in Mexico. Probabilities at **(a)** the first-level PPSBM (all individuals), **(b)** second-level PPSBM (within each first-level cluster; n = 63,196), **(c)** third-level PPSBMs performed on individuals with maximum assignment probabilities to the Central Mexican Highlands (n = 3,408) and, **(d)** fourth-level PPSBMs performed on individuals with maximum assignment probabilities to Central–Isthmus Corridor (n = 124). Gray indicates probabilities of assignment to clusters that were excluded due to extremely small sample sizes (*<* 20 individuals).

Using the templated positional Burrows–Wheeler transform (TPBWT; Freyman et al. 2021), we computed IBD between all participants genotyped on a custom array (∼560,000 SNPs), applying default parameters with the exception of a minimum segment size of 7 cM. For each PPSBM run, we executed 30 independent MCMC chains with 5,000 total iterations and a burn-in of 2,000, using an edge divisor (*z*) of 7 cM and capping the maximum cM shared between close relatives (*β*) at 25 cM. Capping the maximum IBD shared between two individuals in the graph and using a divisor greater than 1 helps reduce biases introduced by close family members and endogamy within the cohort.

We ensured that each independent MCMC chain converged by computing the Geweke Z-statistic and retaining samples from chains with values between −2 and 2 (Geweke 1989). We then removed any cluster that included fewer than 20 individuals with maximum marginal assignment probabilities. With this approach, we identified 9 initial populations across Mexico (Supplementary Table 1, Supplementary Table 2). For each population, we queried all available geographic, language, and cultural survey responses provided by research participants and labeled each population based on the most enriched location, either a broader regional classification or a specific city, county, or state. The labels given to each cluster are mutually exclusive but the geographic representation between clusters may overlap.

To identify sub-structure, we sequentially applied PPSBM clustering to estimate nested hierarchical clusters. Starting from the initial 9 populations identified above, we isolated individuals with the highest marginal probabilities to each population and applied PPSBM clustering, identifying an additional 46 populations (Supplementary Table 1). We subsequently focused on select populations in southern Mexico and continued re-running the PPSBM until sample sizes were too small to detect further structure. This process took us from a set of clusters spanning Southern Mexico (n = 14,838, k = 11), to a set of clusters more narrowly focused in Central Mexican Highlands (n = 3,408, k = 22), and finally, at the most granular level, to a set of clusters in Central–Isthmus Corridor (n = 124, k = 4). Throughout the sub-clustering process, an enrichment of cultural terms became apparent in survey responses; therefore, we labeled certain clusters by these terms if they were enriched. The terminal node of this sub-structuring hierarchy resulted in the identification of four clusters: Córdoba, Isthmus Zapotec speakers in Oaxaca, Lebanese Mexicans, and Jewish Mexicans in Mexico City (Figure 3).

To understand the genetic ancestry background of each identified population, we computed local genetic ancestry estimates using 23andMe’s Ancestry Composition algorithm for each research participant and aggregated the mean genetic ancestry per population (Durand et al. 2021, 2014). The most common genetic ancestry components in this cohort were Spanish & Portuguese (45.0% *±* 15.2%), Indigenous American (41.1% *±* 15.5%), Senegambian & Guinean (1.4% *±* 0.6%), and Ashkenazi Jewish (1.0% *±* 1.8%). However, these genetic ancestry proportions varied among specific populations identified by the clustering procedure. For instance, in the Northern Nuevo León population, mean Spanish and Portuguese ancestry reached its highest value at 62.3%; in the Santiago Zoochila population, Indigenous American ancestry peaked at 86.2%; in the Copala population, Senegambian & Guinean ancestry reached 6.8%; and in the Jewish Diaspora population in Mexico City, Ashkenazi Jewish ancestry reached 29.4% (Figure 4).

**Figure 4:**
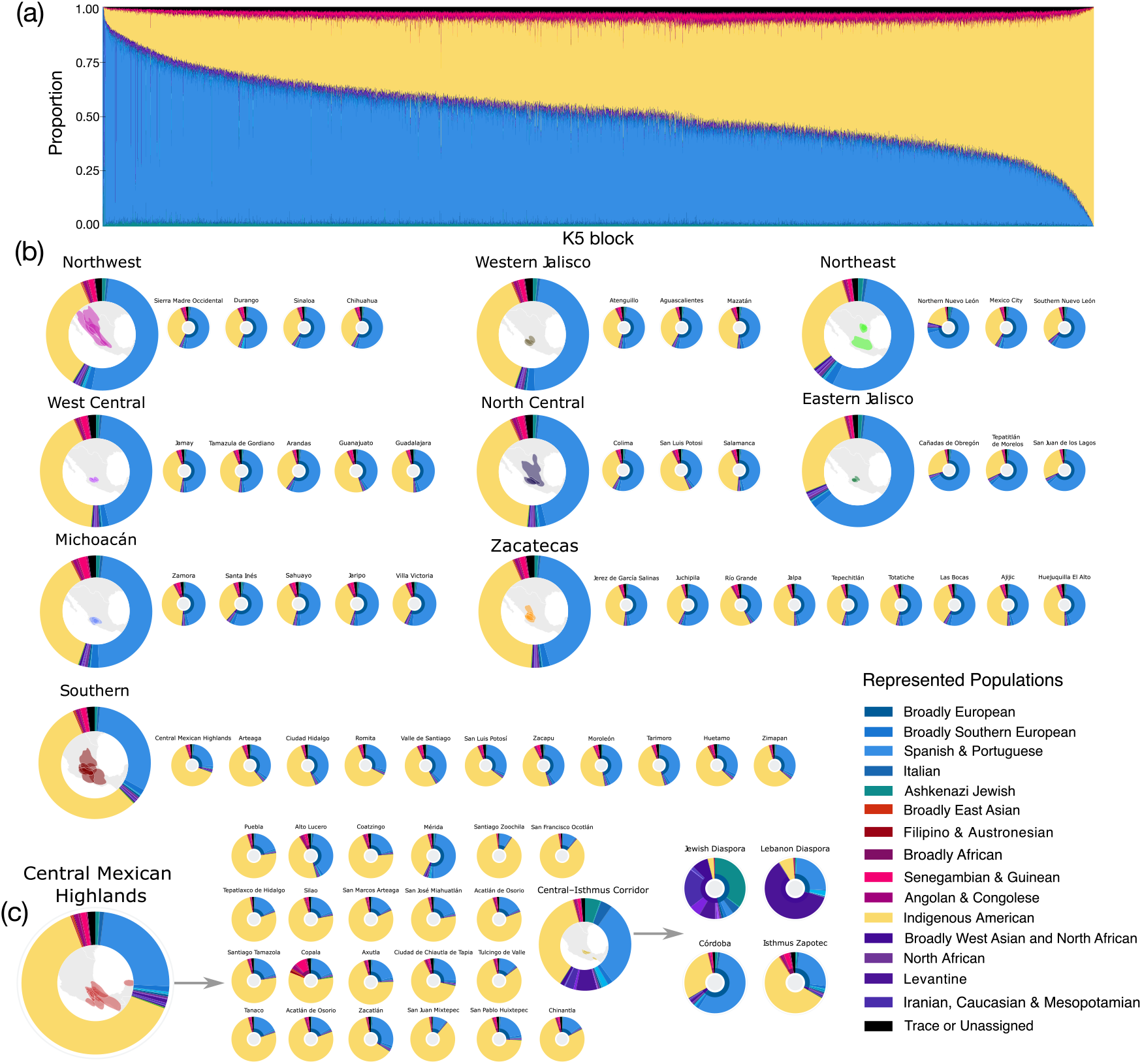
Genetic ancestry proportions of research participants who reported that all four grandparents were born in Mexico. **(a)** Genetic ancestry proportions of individuals in the initial clustering cohort; the x-axis reflects the mean ancestry of every five individuals with the most similar ancestry proportions (for anonymity purposes). **(b)** Mean genetic ancestry proportions of the nine populations identified in the initial round of PPSBM clustering, with the mean proportions of sub-population identified in subsequent sub-clustering shown to the right. **(c)** Genetic ancestry proportions of additional PPSBM clustering applied to the Central Mexican Highlands population, followed by the Central–Isthmus Corridor population. Genetic ancestry populations presented at very low proportions are merged into the “Trace” category. Inner circles of the wheels, when present, represent continental-level ancestry proportions.

We also aggregated IBD results across all populations by calculating the mean total IBD shared between population pairs using the following formula:

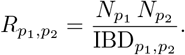

where 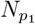 and 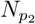 represent the sample sizes of populations *p*_1_ and *p*_2_ and 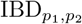 is the total IBD shared between individuals in populations *p*_1_ and *p*_2_. We also computed the mean segment size, using only segments ≥ 8 cM. Populations with the highest within-cluster IBD sharing were those with greater Jewish genetic ancestry, such as Northern Nuevo León, the Jewish Diaspora, and the Lebanon Diaspora. In contrast, populations with the lowest within-cluster IBD sharing tended to be larger clusters such as Southern Mexico and Central Mexican Highlands (Supplementary Table 3).

## 5 Conclusion

By coupling IBD graphs with Bayesian PPSBM clustering, we introduce a principled, uncertainty-aware alternative to heuristic community detection for inferring population structure. Simulations show that the PPSBM recovers fine-scale clusters more accurately than widely used modularity-maximizing approaches, while also flagging recent divergence and admixture through posterior probabilities. Applying the PPSBM to the genomes of 63,196 individuals of Mexican descent demonstrated that this method scales effectively to large genomic cohorts and has the potential to reveal fine-scale structure. Initial PPSBM clustering identified nine broad-scale populations. However, applying PPSBM clustering recursively to these populations—while incorporating local ancestry inference and survey data—uncovered cryptic population structure shaped by more recent historical and demographic events. This combined approach enables the identification of genetic communities shaped by diverse historical processes.

In northern Mexico, some populations—such as those in Nuevo León—exhibited the highest levels of Spanish & Portuguese and North African genetic ancestry, with many individuals identifying as Jewish, suggesting these individuals are likely of Sephardic Jewish descent (Bejarano 2005). This is further supported by elevated within-population IBD, indicative of a founder event (Slatkin 2004). In southern Mexico, further sub-structuring of the Central Mexican Highlands cluster revealed multiple populations shaped by known episodes of migration and admixture. For instance, the Copala cluster, primarily located in Acapulco, included individuals with the highest levels of Filipino & Austronesian as well as Sub-Saharan African genetic ancestry—likely reflecting the influence of the Manila Galleon trade route that connected Manila and Acapulco from the late 16th to early 19th century (Bjork 1998). Sub-structure within the Central–Isthmus Corridor population highlighted multiple diasporic groups, including Lebanese Mexicans and Syrian Jews, who likely migrated to the region in the early 18th century (Hamui-Halabe 1997). Finally, PPSBM clustering identified indigenous communities with very little Spanish & Portuguese admixture, such as the Isthmus Zapotec, concentrated in the Isthmus of Tehuantepec, the Zapotec of Santiago Zoochila, Maya in Mérida, Yucatán, and the Mixtec of San Juan Mixtepec.

Together, these findings demonstrate that the complex history of immigration, colonialism, and slavery in Mexico can be captured and recapitulated using IBD-based PPSBM clustering. We applied recursive sub-structuring to specific populations, resulting in the identification of 81 distinct populations. However, this methodology has the potential to uncover even finer-scale structure given sufficiently large sample sizes. This approach provides a robust foundation for downstream demographic analyses and association studies where subtle population structure is critical, and can inform both genetic ancestry deconvolution and the study of health outcomes associated with specific populations.

## 6 Supplementary Table Captions

**Supplementary Table 1**. *PPSBM subclustering statistics*. Each row corresponds to an independent PPSBM subclustering analysis. Columns include the number of individuals included in the clustering analysis, the final estimate of the number of subclusters, the number of MCMC chains that converged and were retained, the mean estimated number of subclusters (*K*) with standard deviation across all iterations of all MCMCs, the minimum and maximum estimated *K*, the mean Geweke Z-statistic with standard deviation, and the mean modularity across all MCMC iterations.

**Supplementary Table 2**. *Cluster sample sizes and marginal posterior probabilities*. All identified clusters across all PPSBM runs, including their sample sizes and the mean posterior probability of assignment for individuals in each cluster, along with standard deviation.

**Supplementary Table 3**. *IBD sharing statistics within and between all clusters in each PPSBM clustering analysis*. “Mean Total IBD” refers to the total IBD shared (in cM) between two clusters divided by the number of possible pairwise comparisons between individuals. “Mean Segment Size” refers to the average size of IBD segments ≥ 8 cM shared between all individuals. “Within Population” indicates whether the comparison is within a single cluster or between two different clusters.

## Acknowledgements

We thank Tim Do, Nate McQuay, and David Poznik for their suggestions and words of wisdom. We also thank 23andMe customers who consented to participate in research for enabling this study. Finally, we thank employees of 23andMe who contributed to the development of the infrastructure that made this research possible.

